# Character trees from transcriptome data: origin and individuation of morphological characters and the so-called “species signal”

**DOI:** 10.1101/019380

**Authors:** Jacob M. Musser, Günter P. Wagner

**Affiliations:** Yale Systems Biology Institute, West Haven, CT 06516; Yale Peabody Museum of Natural History, New Haven, CT 06511; Department of Ecology and Evolutionary Biology, Yale University, New Haven, CT 06511; Department of Obstetrics Gynecology and Reproductive Sciences, Yale Medical School, New Haven, CT 06511; Department of Obstetrics and Gynecology, Wayne State University, Detroit, MI 48201

## Abstract

We elaborate a framework for investigating the evolutionary history of morphological characters. We argue that morphological character trees generated by phylogenetic analysis of transcriptomes provide a useful tool for identifying causal gene expression differences underlying the development and evolution of morphological characters. They also enable rigorous testing of different models of morphological character evolution and origination, including the hypothesis that characters originate via divergence of repeated ancestral characters. Finally, morphological character trees provide evidence that character transcriptomes undergo concerted evolution. We argue that concerted evolution of transcriptomes can explain the so-called “species-specific clustering” found in several recent comparative transcriptome studies. The species signal is the phenomenon that transcriptomes cluster by species rather than character type, even though the characters are older than the respective species. We suggest that concerted gene expression evolution results from mutations that alter gene regulatory network interactions shared by the characters under comparison. Thus, character trees generated from transcriptomes allow us to investigate the variational independence, or individuation, of morphological characters at the level of genetic programs.

## Introduction

In evolutionary genetics and molecular evolution there is a tendency to avoid addressing the evolution of complex traits. There are many good reasons for this, but perhaps the primary obstacle has been the lack of tools useful for addressing the experimental challenges inherent in the subject. This is particularly true for morphological, physiological, and behavioral traits, which occupy a gray zone of inherent importance but limited mechanistic knowledge. These complex phenotypic traits share the feature that there is no clearly understood relationship between genetic information at the DNA level and their physical organization at the phenotypic level. Further, these traits must be re-created in each generation, and thus their pattern of inheritance is quite complex, even to the point where one may question whether it is possible to speak of the heritability of complex phenotypic traits at all. Nevertheless these traits are regenerated every generation quite reliably, remain recognizable across hundreds of millions of years of evolutionary history (=homology; Wagner, 2014), and exhibit quasi-independence that enables distinct variational properties and the evolution of different functional roles.

Recent advances in functional genomics have fundamentally changed the landscape for comparative biology and the study of character evolution (Roux et al., 2015). In particular, RNAseq can be used to estimate expression of all genes or transcripts annotated in a species (transcriptomes) for cells, tissues, and organs (Ozsolak and Milos, 2010). Transcriptomes produced via RNAseq represent more than simply an aggregation of expression values, but rather offer a system-wide view of genomic activity at a given place and time in the organism. Here we review an analytical framework that takes advantage of RNAseq to study the evolution of cells, tissues, and organs: the use of transcriptomes for building phylogenetic trees that depict hypotheses of historical relationships among morphological characters and for identifying candidate gene expression changes related to character origination.

Phylogenetic methods have traditionally been used to reconstruct evolutionary relationships among genes and species. More recently, researchers in the field of developmental evolution (devo-evo) have applied these methods to the study of phenotypic characters at a variety of organismal scales (Serb and Oakley, 2005). This includes multi-cellular morphological structures (Coates et al., 2002; Geeta, 2003; Oakley, 2003; Wang et al., 2012; Tschopp et al., 2014; Musser et al., 2015), cell types (Arendt, 2003; 2008), developmental fields (Liu and Friedrich, 2004), metabolic pathways (Cunchillos and Lecointre, 2005), and protein networks (Plachetzki et al., 2007). In these studies, phenotypic characters, instead of genes or species, are treated as operational taxonomic units (OTUs) and placed at the tips of the tree. The process by which these trees are generated typically involves deconstructing phenotypic characters into component parts, coding character states for these components into a data matrix, and then using this data matrix to construct a tree using common phylogenetic methods (e.g. parsimony or maximum-likelihood approaches). For instance, Coates et al. (2002) coded states for morphological characteristics shared by paired appendages in a number of stem tetrapods. Coates et al. (2002) then conducted a parsimony analysis with this dataset to look for evidence of correlated changes in morphology during the early evolution of the tetrapod fore- and hindlimb.

Transcriptomes provide a standardized method of comparison for investigating the evolution of cell, tissue, and organ *types* (hereafter referred to as morphological characters). Transcriptomes are particularly useful for evolutionary studies of morphological characters for multiple reasons. First, as all cells within an organism share the same genome (ignoring somatic mutations and programmed gene re-arrangements like in adaptive immunity), in principle it is possible to quantify and compare expression for all genes/transcripts among morphological characters. Second, transcriptomes provide a “snapshot” of the activity of the underlying gene regulatory network (i.e. developmental program). Thus, transcriptomes are useful in that they allow phenotypic characters to be studied through the activities and evolution of their gene regulatory networks.

Here, we argue that using transcriptomes to reconstruct hierarchical, historical relationships among morphological characters is useful for:

- Investigating the origin and homology of morphological characters
- Identifying causal gene expression changes for novel morphological characters
- Measuring the degree of individuation among characters using the so-called “species signal”
- Testing different models of morphological character evolution

These last two uses investigate two concepts in devo-evo: (1) that new phenotypic characters arise via the divergence of repeated ancestral characters (i.e. they display a tree-like structure of relationships; Riedl, ′78; Muller and Wagner, ′91; Oakley, 2003; Serb and Oakley, 2005; Arendt, 2008, Liang et al., 2015, Kin et al., 2015), and (2) that phenotypic characters originate by becoming evolutionarily/variationally individuated (Wagner, 2014). Here we use the term “individuated character” to mean a morphological character that evolves independently of other morphological characters in the sense of Lewontin’s notion of genetic quasi-independence (Lewontin, ‘78). This notion of evolutionary individuation emphasizes the ability of characters to vary to some degree independently (i.e. their variability), rather than simply the presence of some given phenotype difference. In individuated morphological characters, mutations may cause phenotypic changes in one character but affect other characters less or not at all. In contrast, characters that are not individuated exhibit the same phenotypic responses to mutations. At the level of gene expression, characters lack individuation when sharing all or most of the same regulatory connections between specific regulatory molecules (e.g. transcription factors or transcription factor complexes) and cis-regulatory elements. A mutation that alters an interaction between a regulatory molecule or complex with a cis-regulatory element results in correlated gene expression changes in all characters employing the regulatory interaction. The degree of individuality can then be understood as the degree of overlap of regulatory interactions shared by two characters. Characters that share less overlap of gene regulatory network connections exhibit a higher degree of individuality.

Incomplete individuation has consequences for the interpretation of character trees generated from transcriptomes. Most hierarchical clustering algorithms and phylogenetic reconstruction methods assume that a gene’s expression changes independently in different characters. However, if characters are not fully individuated, due to the sharing of gene regulatory network interactions, then the assumption of independence among these characters is violated. Evolutionary changes in gene expression that occur simultaneously in multiple characters will thus be treated as synapomorphies (shared derived traits) leading to the erroneous conclusion that different characters within a species are more closely related to each other than to the homologous characters in a related species. We think this is the reason for the species-specific clustering frequently observed in transcriptomic data (Lin et al., 2014; Pankey et al., 2014; Tschopp et al., 2014; Yue et al., 2014). Furthermore, we think that the degree to which characters are individuated influences their propensity to exhibit species-specific clustering. Thus, the topology of character trees generated from transcriptome data reflects the interplay of the historical relationships among morphological characters, as well as the degree of evolutionary individuation of those characters.

In the following sections we first discuss existing evidence for the idea that phenotypic characters, like genes and species, may be treated as historical individuals that share common descent. As part of this we discuss the connection between serial homology, individuation, and common descent of morphological characters. We also highlight an application of a morphological character tree from the literature, i.e. cell type trees (Arendt, 2008). In the subsequent section we discuss the interpretation of character trees generated from transcriptome data. We show how character trees generated from transcriptome data may be used to identify historical relationships among morphological characters, and to identify candidate causal gene expression changes underlying the origin of novel characters. In the final section we discuss technical issues important to consider when analyzing transcriptomes.

## Serial homology, common descent, and individuation of morphological characters

A common feature of multicellular organisms is that they are composed in large part of repetitions of the same or similar building blocks (Owen, 1848; Riedl, ′78; Remane, ′56; Raff, ′96; Weiss, ′90). Examples are the multitude of identical or similar cells of most cell types, tissue types such as smooth or skeletal muscles, and more complex structures such as the leaves on a plant, bird feathers, or segments of annelid worms. These repetitive units exhibit two notable features. First, although variation among different morphological characters exists, there is often not a continuum of intermediate variation (e.g. among cell types; Vickaryous and Hall, 2006). Thus, morphological characters exist as semi-discrete units. This was recognized at least as early as Waddington (‘57), and was the inspiration for his concept of “canalization”. The second notable feature of repeated morphological characters is they exhibit at least partial context insensitivity (e.g. are developmental modules; Wagner, ’96; Raff, ‘96). By this we mean multiple instances of the same morphological characters maintain their identity despite existing in different spatiotemporal locations and environments within the organism, and also may be experimentally induced in ectopic locations (e.g. eyes on the wing and legs of Drosophila; Halder et al., 1995).

Early anatomists recognized this repetition of similar parts as another dimension of the homology concept. Owen (1848) coined the term serial homology, the same structure repeated within the same organism, as a complement to “special homology”, the presence of the same organ in different species. With the advent of the phylogenetic interpretation of interspecific or special homology by Lankester (1870), however, the complementarity between special and serial homology was compromised. If homology is explained by descent from a common ancestor, “serial homology” cannot be homology at all (Remane, ’56; Patterson ‘82). The treatment of serial homology as distinct from special homology served a useful purpose in defining tests for special homology. For instance, Patterson (1982) proposed the conjunction test, which failed if proposed homologues were found to occur within the same organism. However, treating serial homology as distinct from special homology clouded recognition of the shared mechanistic bases between the two forms of homology.

Complementarity of special and serial homology can be restored, at least to some degree, if morphological characters, both within and among species, are recognized as historical entities that exhibit distinct evolutionary histories. Morphological characters exist as historical entities because multicellular organisms have evolved compartmentalization of developmental pathways. Stated more simply, organisms are composed of parts, each with their own gene expression program, which enables that character to vary quasi-independently from other characters. In turn, variational quasi-independence allows these characters to have their own evolutionary history and evolve their own functional specializations. Several recent papers have proposed that the mechanistic basis for different characters may be the differential deployment of a specific set of core regulatory genes, described in various ways as character identity networks (ChIN; Wagner, 2007), kernels (Erwin and Davidson, 2009), or core networks (Graf and Enver, 2009). These concepts highlight that the individuality of body parts and cell types is tied to highly conserved parts of the gene regulatory network. These core genes play the specific functional role of enabling differential gene expression, necessary for the developmental and physiological phenotype of a character. In other words, the character identity network associated with a character explains how that character *can* be, but does not have to be, phenotypically distinct from other characters in the same organism. The character identity network also explains how characters may vary quasi-independently from each other.

Special homologs are the result of the deployment of the same character identity network in different organisms. Serial homologues are the result of deploying the same, or historically related, character identity networks in different locations of the body or different individuals of the same species (sexual dimorphisms and complex life cycles). In contrast to special homology, the degree of difference among the implemented gene regulatory networks within the same organism depends on how individuated the characters are. Structures that exhibit no individuation, for instance two identical and adjacent hairs, may share identical core networks and identical or highly similar overall gene regulatory networks. On the other hand, forelimbs and hindlimbs exhibit a larger degree of individuation (Naiche et al., 2005; Duboc and Logan, 2011) and have evolved at least partially distinct character identity networks enabling gene expression to evolve quasi-independently during development (Ouimette et al., 2010; Taher et al., 2011). Hence, from a mechanistic point of view, special and serial homology can be seen as reflecting the same fundamental fact, *namely that body part identity and development are controlled by conserved elements within gene regulatory networks and that body parts in different organisms, or in different locations of the same organism, are “the same” because they result from the implementation of historically related character identity networks*. For that reason both special and serial homologs are legitimate and complementary objects of study.

If morphological characters are historical entities, how do they originate? Given that many aspects of morphology are repetitive, with varying degrees of evolutionary individuation, new characters may arise via the divergence of repeated ancestral characters and their character identity networks. The result is a hierarchical pattern of common descent among morphological characters (Riedl, ′78; Muller and Wagner, ′91; Raff, ′96; Oakley, 2003; Arendt, 2008; Oakley and Rivera, 2008). An analogy may be made to gene duplication and divergence. However, there are several important differences between genes and morphological characters. These differences arise because genes reproduce directly by semiconservative replication, whereas morphological characters are recurring structures that must be rebuilt every generation. The consequence is that gene duplication can result in the immediate individuality of gene paralogs, with subsequent mutations solely affecting one copy or the other. In contrast, multiple instances of a morphological character are the result of duplications occurring every generation by deployment of the same gene regulatory network in different spatiotemporal locations during development. For morphological characters, only the instructions for the gene regulatory network and its deployment, embedded within the genome, are inherited directly. Thus, multiple instances of a morphological character may not exhibit any individuality despite existing at different spatial or temporal locations within the organism. Morphological characters that lack individuation undergo concerted gene expression evolution, as most mutations that affect the shared gene regulatory network will result in gene expression changes across multiple instances of the morphological character in the individual. For the purpose of depicting historical relationships among morphological characters, multiple instances of a morphological character present within the same individual should be represented as each other’s closest relatives.

Interestingly, there is a similar example of concerted evolution among duplicated genes. This is the case of ribosomal RNA genes, where gene trees that included paralogs sampled across multiple species found that paralogs from the same species were each other’s closest relatives, despite the ancient origin of this gene family (Dover and Coen, ′81; Coen et al., ′82). This concerted evolution in ribosomal RNA genes is the result of unequal crossing over and gene conversion. Thus, although the mechanism for concerted evolution (or lack of individuation) of ribosomal RNA genes differs from that of morphological characters, it results in a common phylogenetic pattern where copies of genes or characters within the same species are each other’s closest relatives.

In contrast to most gene paralogs, morphological characters may vary substantially in the degree to which they are individuated. To distinguish between non-individuated and individuated characters, we use the terms homomorph and paramorph respectively (Wagner, 2014). Homomorph characters are those that are identical copies, and share identical character identity networks and very similar gene regulatory networks overall. In terms of gene expression they may only differ with respect to the genomic information necessary for specifying the spatiotemporal location of their respective deployments.

Paramorph characters are individuated morphological characters related via descent and differentiation from an ancestral character, and exhibit distinct character identity networks and gene regulatory states. A large fraction of real characters, however, are likely somewhere in-between the two extremes of homo-(i.e. identical) and paramorph (i.e. fully individuated) characters. They have acquired some degree of evolutionary individuality but still are highly likely to share derived features because of the similarity of their gene regulatory networks. Nevertheless, partially individuated characters can exhibit sufficient unique variability such that they are able to adapt to different selection pressures. A prime example are the various skin appendages of amniotes, in particular the functionally differentiated kinds of feathers on a flying bird (see chapter 9 in Wagner 2014 for more detail). If a group of characters remains only partially individuated over longer periods of evolution they can be called a “character swarm,” in analogy to the notion of a species swarm, which is a group of partially isolated populations (Gentry, ′82).

If new morphological characters (paramorphs) arise via divergence of repeated ancestral characters, we should expect to find evidence of this hierarchy and history of common descent. One example of where this has been documented is in the evolution of metazoan photoreceptor cells (Fig. 1; Arendt, 2008). Bilaterians exhibit two distinct classes of photoreceptor cells, ciliary and rhabdomeric. By coding various attributes of photoreceptors cells, including presence/absence of different opsins, physiological pathways, and subcellular structures, Arendt (2008) generated a tree depicting evolutionary relationships among photoreceptor cell types. Under this model, termed the sister cell type model, new photoreceptor cell types arose via divergence of multifunctional ancestral photoreceptor cell types. This occurred both via segregation of ancestral molecular functions in the descendent cell types, as well by the evolution of new functions. An interesting finding of this model is that the rhabdomeric photoreceptor cell lineage, which serves a primarily visual function across most animals, is present in vertebrates but instead serves a non-visual function in the retina.

**Figure 1:**
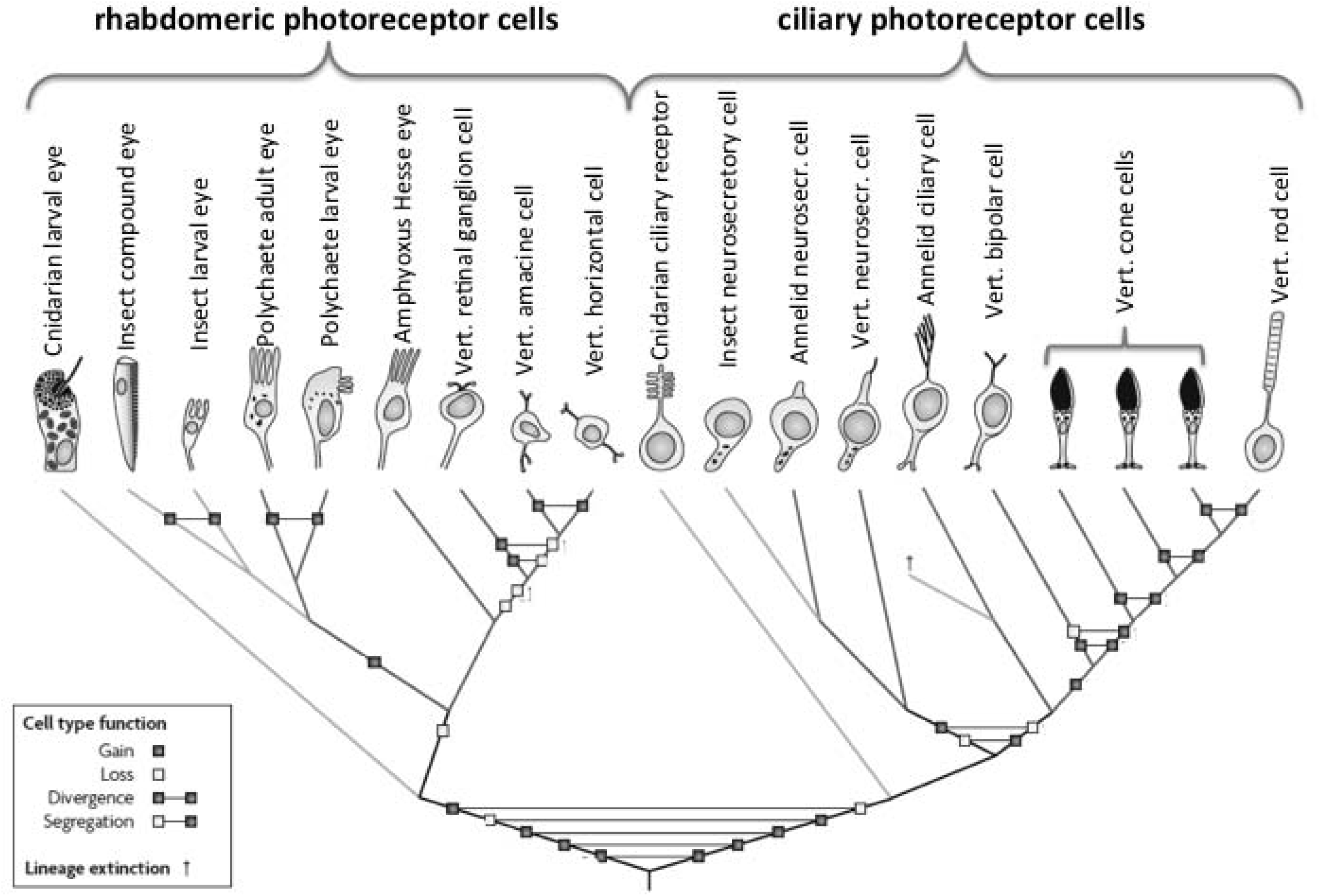
Cell type tree of metazoan photoreceptor cell types (after Arendt, 2008 with permission of Nature Publishing Group). Photoreceptor cells come in two major clades, rhabdomeric and ciliary photoreceptor cells. Notably, certain vertebrate retinal interneurons are closely related to rhabdomeric photoreceptors, and likely evolved from an ancestral rhabdomeric photoreceptor cell type. Internal nodes in the tree represent ancestral cell types. Branching events represent either the origin of a cell type by bifurcation of an ancestral cell type, or speciation events. Shading of lineages indicate taxa or groups of taxa.

## Interpreting Character Trees generated from transcriptome data

In this section we discuss how character trees generated from transcriptome data may be interpreted and used to investigate character evolution. Although we reserve a more technical discussion of transcriptome analysis for the final section, it is worth highlighting a few principal points here. First, as mentioned previously, morphological character trees treat morphological characters (e.g. cell, tissue, and organ types) as operational taxonomic units, which are placed at the tips of the tree. Generating these trees from transcriptome data requires applying phylogenetic methods, for instance parsimony or maximum likelihood, using a matrix composed of gene expression values. This matrix is analogous to an alignment of DNA sequence data used to generate phylogenies reflecting historical relationships among species. However, whereas a matrix of DNA sequence data encodes information about different base pairs among different species (base pair by species), a matrix of gene expression data is composed of information regarding the expression of different genes among different morphological characters (gene by morphological character). Analyzing these matrices will ultimately require mathematical models of gene expression evolution.

### Relationships of the same morphological character across species

Character trees generated using transcriptomes collected from the *same* morphological character in *different* species are useful for identifying species-specific gene expression changes in the given morphological character. Although generation of the species tree itself is possible using gene expression data, if the species tree is known it may be worthwhile to constrain its topology, and then map expression changes on it. If gene expression is treated as a binary character (Wagner et al. 2013), where genes are called expressed or not expressed, it is also possible to utilize a variety of ancestral state reconstruction tools to infer expression changes occurring on ancestral branches. This may be useful for identifying large-scale gene expression gain or loss events in the history of a character, and correlating expression change events with functional evolution in the character.

### Relationships of different morphological characters within a species

Another type of character tree is one composed of *different* morphological characters that are present within the *same* organism. This type of character tree is a phylogenetic hypothesis of historical relationships among serially homologous morphological characters. It provides a historical framework useful for testing hypotheses of relatedness among different morphological characters. It also places the evolution of gene expression within a historical context, which allows for distinguishing derived gene expression differences associated with a particular character, from ancestral changes resulting in expression shared by a clade of related characters (Fig. 2). Failure to make this distinction may lead to misleading statements regarding character evolution. For instance, gene expression in feather development has now been documented for a large number of genes (c.f. Lin et al., 2006 for a review). Many of these genes are expressed early in feather development at or shortly after the placode stage, when the only anatomical structure present is a thickened epidermis and small bump on the skin. A recent study by Lowe et al. (2015) investigated feather evolution by identifying the branch of the vertebrate tree along which “feather” genes and associated conserved non-coding elements originated. They found the highest rate of origin for conserved non-coding elements was prior to the amniote ancestor, long before feathers evolved. However, there is mounting evidence that many of these genes, which are expressed in the feather placode, are also expressed in similar patterns in avian feet scale placodes (Crowe and Niswander, ′98; Widelitz et al., 2000; Yasue et al., 2001; Harris et al., 2002; Tao et al., 2002; Lowe et al., 2015; Musser et al., 2015) and even in the hair placodes of mammals (St-Jacques et al., ′98; DasGupta and Fuchs, ′99). Although we still lack a character tree of feathers and other skin appendages generated with transcriptome data, a preliminary character tree hypothesis (Musser et al., 2015) suggests that regulatory molecules important in feather placode development evolved in an ancestral skin appendage prior to the amniote ancestor. Thus, when studying this stage of development it may be more appropriate to recognize these genes as “placode” genes or even “amniote skin appendage” genes, rather than “feather” genes. If this inference is correct, it explains the finding of Lowe et al. (2015) that the regulatory elements of so-called “feather” genes originated prior to the most recent common ancestor of amniotes rather than in the stem lineage of birds, where true feathers actually evolved.

**Fig. 2:**
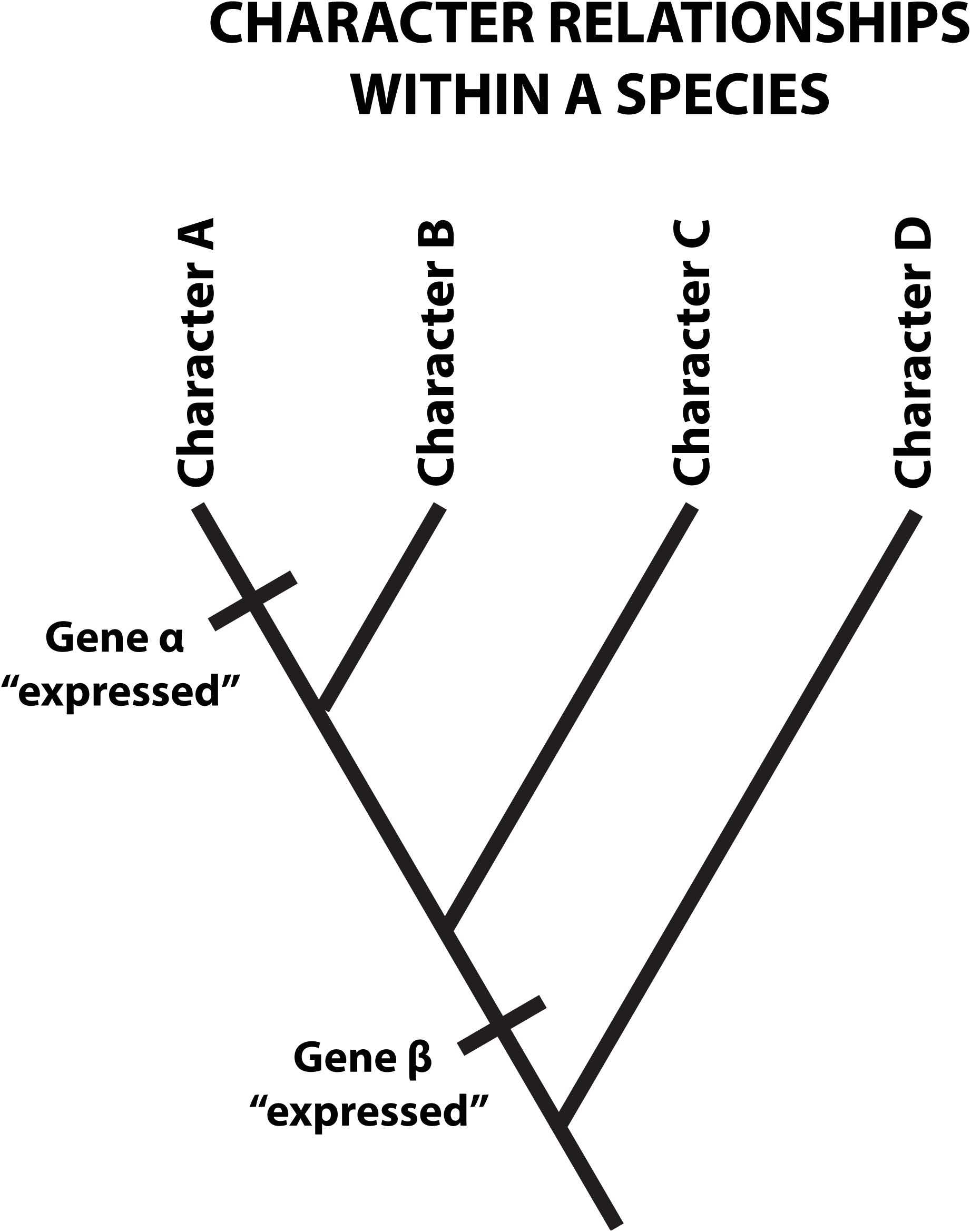
Relationship of serially homologous characters within a species. The tree depicts interrelationships among four characters. Expression of gene β evolved in an early ancestor and is unlikely to explain traits specific to character A. Expression of gene α evolved on the branch leading to character A, and is a candidate for explaining traits specific to character A.

The distinction between placode genes and feather genes is not only of semantic value since it raises the question of what the expression of these genes actually explains. If we call a gene, or a gene regulatory network, a “feather” gene or network, we imply that we have discovered the genetic information for making or evolving a feather. With respect to the “feather” placode genes, that may be mistaken, since these genes in fact likely explain the development of an embryonic feature that is shared among multiple skin appendages, including both feathers and feet scales in birds, and thus do not teach us how feathers develop, or how they evolved.

In a recent example, Kin et al. (2015) generated a character tree of mesenchymal cell types to test alternative hypotheses of the origin of the endometrial stromal cell (ESC), as well as identify regulatory molecules causally relevant to ESC evolution. Endometrial stromal cells are a type of mesenchymal cell found in the uterus of mammals, and are important in regulating the mother’s response to the placenta during pregnancy (Gellersen and Brosens, 2014). Previous evolutionary hypotheses have suggested endometrial stromal cells might be related to either myofibroblasts or follicular dendritic cells (Oliver et al., 1999; Dunn et al., 2003; Munoz-Fernández et al., 2006). Kin et al. (2015) found that endometrial stromal cells and follicular dendritic cells formed a well-supported clade, whereas a close relationship of endometrial stromal cells and myofibroblasts was rejected. They then performed an ancestral state reconstruction to identify gene expression changes associated with the evolution of endometrial stomal cells. Using RNAi-mediated knockdowns, Kin et al. (2015) found that transcription factors gained on the most recent branch leading to endometrial stromal cells were more likely to regulate endometrial stromal cell marker genes than were transcription factors gained on earlier branches leading to ancestors shared between endometrial stromal cells and other mesenchymal cell types.

Two other recent studies provide additional examples of how character trees depicting relationships of characters within the same organism may be useful for investigating character evolution. Wang et al. (2012) assayed transcriptomes from developing digits of the chick embryonic forelimb and hindlimb to test the hypothesis that the avian digit growing out of the 2^nd^ (anterior-posterior) position in the developing avian hand (aka “wing”) was actually the thumb, which had undergone a frameshift and moved to position two from an ancestral digit position one (see Wagner and Gauthier, ‘99 for background). The resolution of this debate has important implications for the hypothesis that extant birds are descended from dinosaurs (c.f. Prum, 2002; Young et al., 2011; Xu et al., 2014 for reviews of this debate). Hierarchical clustering of forelimb and hindlimb digits using transcriptomes revealed the digit growing from position 2 of the avian hand is most closely related to the digit growing from position 1 in the avian foot. This confirmed its identity as the thumb, and was consistent with both the frameshift hypothesis and dinosaurian origin of birds (Wagner and Gauthier, ′99). In another, more recent example, Tschopp et al. (2014) demonstrated a close relationship between limb buds and external genital buds, relative to tail buds, in mouse, chicken, and several squamates. In the lizard *Anolis carolinensis* genital buds (hemipenes) grow from the hindlimb, and hierarchical clustering of gene expression from these structures resulted in a well-supported limb and genital bud clade, with tail buds more distantly related. Furthermore, early genital buds were embedded within limb buds, and were more closely related to hindlimb buds than forelimb buds. In mouse, the tree topology differed, with early genital buds sister to fore- and hindlimbs, although they still formed a well-supported genital and limb bud clade with tail buds more distantly related. The tree topology in mouse is notable because the mouse genital bud develops from the base of the tail bud. Thus, although there is a difference as to whether early genital bud development in mouse and lizard is embedded within limbs, in both cases there is a closer relationship between limb and genital buds than either tissue has with tail buds, suggesting the character tree does not simply track developmental position. There are several possible evolutionary interpretations of the character trees in Tschopp et al. (2014). The first possibility is that early genital buds and early limb buds are serially homologous structures, and share related character identity networks. This explanation in no way implies that later limb and genital structures are closely related, but rather recognizes that complex structures such as limbs and genitalia may be serially homologous in early development, but acquire distinct identities at later stages of development. An alternative to the hypothesis of serial homology is that genital buds independently co-opted expression of genes used in limb bud development. In this case, the regulatory connections in genital and limb bud regulatory networks are expected to be different, despite yielding similar expression patterns of some genes, which drives the clustering pattern.

### Relationships of different morphological characters across species

The last type of character trees we will discuss are those that include *different* characters sampled across *multiple* species (Fig. 3). This type of morphological character tree is useful for addressing the questions discussed in the previous sections, but also allows for testing whether different characters are evolutionarily individuated, or whether they are undergoing concerted gene expression evolution with respect to each other. Individuated characters are those that may evolve independently, with mutations that tend to affect gene expression in one character but not others. In contrast, in characters that lack individuation due to shared use of the same gene regulatory network interactions, mutations will tend to affect gene expression in both or multiple characters simultaneously. Characters that evolve independently, or mostly independently, will be expected to cluster by homologous characters, rather than by species (Fig. 3). That is so because the transcriptome of each character will accumulate expression changes independently, and clustering will reflect the historical signal of homology that unites the same character across different species. Related characters undergoing concerted evolution will cluster by species because mutations altering shared genetic machinery will result in similar changes in gene expression across the characters. When these changes occur after the speciation event for the species under comparison, these mutations will generate species-specific synapomorphies. Given enough of these mutations, this signal may override the historical signal of homology, and result in clustering of character transcriptomes by species.

**Fig. 3:**
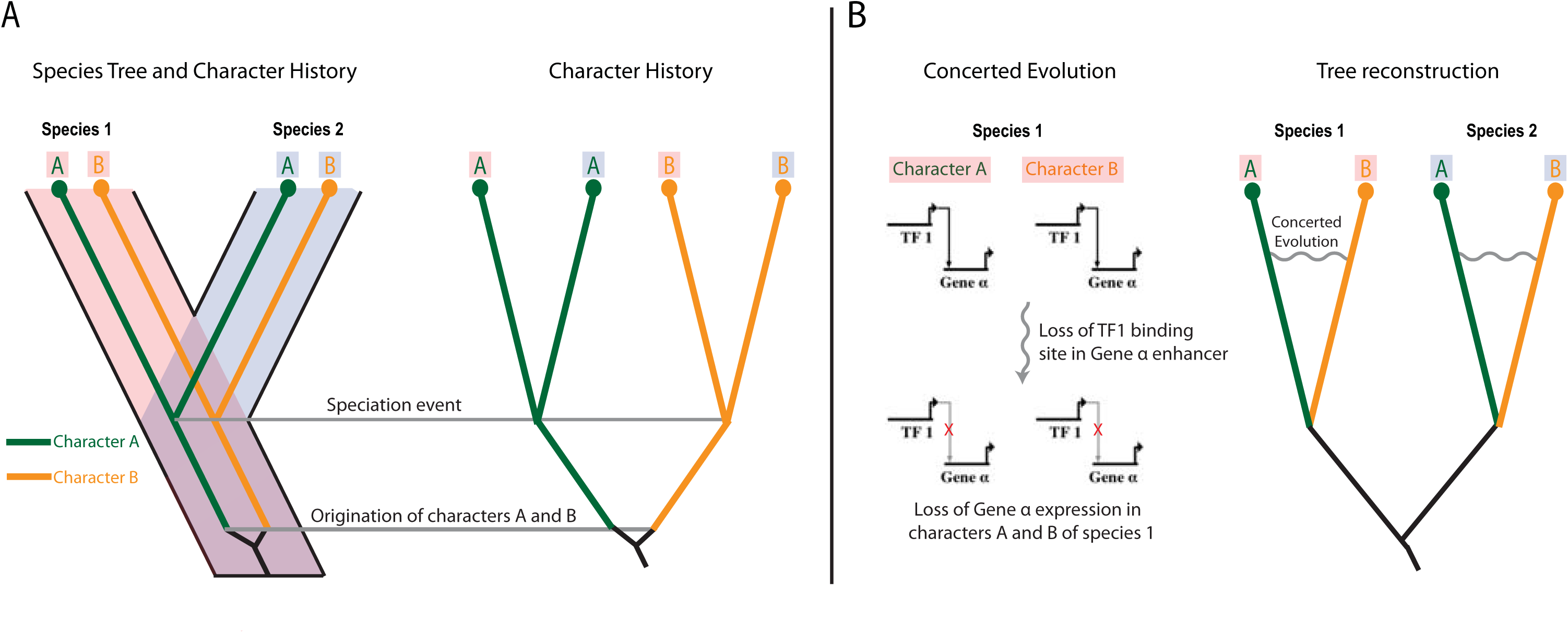
Concerted evolution influences character tree reconstruction. A) Species tree and true character history. Characters A and B originated in an ancestor prior to the divergence of species 1 and 2. B) Concerted gene expression evolution in characters A and B of species 1. Loss of a shared TF binding site results in loss of gene α expression in characters A and B in species 1, producing a synapomorphy that supports clustering by species.

Two recent papers highlight these alternative examples of characters evolving in an individuated fashion versus concerted evolution. Merkin et al. (2012) generated transcriptomes from various mature tissues, including lung, muscle, testes, brain, and others. These tissues were sampled from four species of mammals and chicken. Hierarchical clustering of gene expression found most transcriptomes clustered by tissue, although some tissues remain clustered by species. This suggests that most of the sampled tissues are strongly individuated at the level of both character identity and gene regulatory network, as gene expression changes in one tissue do not generate similar gene expression changes in other sampled tissues from that species (i.e. they do not generate spurious “species synapomorphies” among the sampled characters). Further, the gene expression identity which unites the tissue types, and which likely evolved prior to the amniote ancestor (as only amniotes tissues are sampled), has been preserved despite millions of years of evolutionary divergence among these species.

In contrast to the expectation that homologous characters cluster together, in their study on the evolution and development of amniote genitalia, Tschopp et al. (2014) found that the first principal component of gene expression variation in a principal components analysis encoded species identity, and also that a tree generated with all sampled transcriptomes resulted in clustering by species, rather than tissue type (Patrick Tschopp, pers. comm.). There are two possible explanations for the pattern of characters clustering by species. First, the characters sampled may have originated more recently than the common ancestor of the species. In the case of fore and hindlimbs of mice and lizards this is not the case. Alternatively, clustering by species is expected if gene expression is evolving in a concerted or correlated fashion among the characters in each lineage, which would be expected if the sampled characters are serial homologs. Concerted gene expression occurs when mutations alter gene expression simultaneously in multiple characters. In a phylogenetic reconstruction these expression changes are interpreted as derived states that are shared by characters within a species, and thus leads to a tree in which the characters are sorted by species rather than historical origin. For that reason when there is a clear “species signal”, the amount and the composition of this signal can be interpreted as evidence of concerted transcriptome evolution, and thus as a direct measure of the degree of variational individuality, or lack thereof. At this point, it is not immediately apparent the extent to which characters must be individuated before they are likely to cluster by species. To determine this, and to extract a quantitative measure of individuation from this type of data, requires a mathematical model of transcriptome evolution (Liang, Musser and Wagner, in preparation).

In cases of incomplete evolutionary individuation the structure of the character tree may also be influenced by the developmental lineages of the characters. As acknowledged earlier, even in the case of homomorph characters the genome must encode information that deploys the same character identity network in different spatiotemporal locations, and differences in spatiotemporal location will be reflected in the transcriptome. In Tschopp et al. (2014), even though the genital buds are more closely related to limb buds than to the tail bud in all sampled species, the genital buds are embedded within limbs in lizards. As the authors note, this is likely due to the developmental origin of the lizard genitalia (the two hemipenes) on the hindlimb buds. Thus, in cases where characters are not fully individuated, for instance as may be the case in early limb and genital bud development, gene expression patterns reflecting positional information may affect the structure of the character tree.

A recent online discussion^1^, with detailed analysis to be published later (Gilad and Mizrahi-Man, forthcoming), has suggested that “batch” effects may be the cause of transcriptomes clustering by tissues in recent studies by the mouse ENCODE consortium (Yue et al., 2014; Lin et al., 2014). Investigation of the mouse ENCODE consortium dataset revealed that most human and mouse transcriptomes were sequenced on different sequencing machines and/or flow cell lanes, potentially confounding the results. This experimental design makes it difficult to distinguish between true biological gene expression differences and batch effects caused by sequencing machine or flow cell lane. However, a close examination of the mouse ENCODE consortium experimental design reveals that human and mouse brain tissue were sequenced on the same lane. Thus, if batch effects were the principal reason for clustering by species, it would be expected that at least brain tissue from mouse and human would cluster together. However, this is not what is observed. Rather, mouse and human brain tissue each clusters with other tissues from their respective species. Thus, although it is important to implement good experimental design in any transcriptomics study, batch effects due to sequencing machine or lane are likely not the principal reason for transcriptomes clustering by species in the mouse ENCODE consortium studies. Furthermore, Lin et al. (2014) observed that some tissues, for instance testes, exhibited a greater degree of tissue-specific gene expression. Tissues with more tissue-specific gene expression were preferentially sampled in comparative studies that found clustering by tissue (Merkin et al. 2012). This observation is consistent with our explanation that greater evolutionary individuation facilitates the evolution of greater tissue-specific gene expression. Thus, the finding by Merkin et al. (2012) that human and mouse transcriptomes cluster mostly by tissue is explained by the fact that they sampled tissues with a high degree of evolutionary individuation with respect to each other.

### Testing alternative models of character evolution

A central prediction of the model of morphological character evolution discussed here (e.g. the sister cell type model in Arendt, 2008 or replication and divergence of more complex morphological characters in Oakley, 2003) is that morphological characters within an organism are related via a hierarchical pattern of common descent. However, an alternative model can be imagined in which new morphological characters arise by combining parts of gene regulatory networks that evolved in unrelated characters (e.g. Niwa et al., 2010; Clark-Hachtel et al., 2013). We call this the hybridization model of character evolution, as new characters arise via the combination of unrelated gene regulatory networks. Under the hybridization model of morphological character evolution, relationships among different morphological characters are better represented as a network, rather than a tree, due to the reticulation caused by gene regulatory network “hybridization” events.

To test these alternative evolutionary models of morphological character origination, Liang et al. (2015) developed a statistical model for the comparison of transcriptomes that predicts the probability of observing a given level of “treeness” by chance. This model thus allows one to calculate a p-value for rejecting “treeness” due to chance, and thus testing the hypothesis that the data analyzed in fact has tree structure. The method was then applied to cell type transcriptomes from human and mouse and found that the transcriptomes of normal cell types have substantial amounts of tree structure. This finding is consistent with the sister cell type model (Arendt, 2008), and more generally with the idea of common descent and divergence of morphological characters discussed in this paper. However, although Liang et al. (2015) found evidence of significant tree structure among the transcriptomes of different cell types, not all comparisons were significantly tree-like. There are several possible explanations for this. One possibility is that measures of “treeness” are low if there is a large amount of gene expression homoplasy. The “treeness” statistic of Liang et al. (2015) is quantified for quartets of taxa, which may be particularly susceptible to homoplasy when the cell types being compared are distantly related. Another possible explanation for lack of “treeness” in some cases may be that certain cell types do indeed arise via the hybridization of distinct gene regulatory networks. To disentangle these two possibilities requires determining whether patterns of gene expression contributing to a lack of “treeness” are the result of similar or different gene regulatory networks in the quartet of cells being compared. If a lack of “treeness” is the result of homoplasy in gene expression, we would expect this to be driven by different underlying gene regulatory networks. On the other hand, the hybridization model of cell type evolution would predict that similar patterns of gene expression that result in a lack of “treeness” are the result of similar gene regulatory networks. Thus, additional knowledge of the actual gene regulatory networks themselves, beyond simply gene expression, will be necessary to determine whether lack of “treeness” arises from a hybridization model of evolution versus homoplasy. In addition to these two possibilities, lack of a tree-like structure may also arise due to a lack of individuation among cells. This would be expected in cases of homomorphs, for instance, where sampled characters are nearly identical, and variation among them is not distributed hierarchically. This possibility was recognized by Liang et al. (2015), and was used to test whether it was appropriate to treat two cells sampled from different parts of the body as distinct cell types. This was done by testing for “treeness” using a quartet composed of two replicates each of two cell sample types. Certain cell types, including a number of different fibroblasts, lacked tree structure, suggesting they are better thought of as the same cell type sampled from different locations in the body.

## Technical Issues when using Transcriptomes to Build Character Trees

In this section we briefly introduce a number of technical issues that the investigator has to confront when analyzing transcriptomes to trace the evolutionary history of cell types and other morphological characters. Many of the issues have not been fully settled and are discussed here in order to facilitate debate.

### Quantification, transformation, and comparison

High-throughput RNA sequencing data typically represent the relative abundance of different RNA species in a sample. This data can be transformed in various ways into expression estimates, where the most parsimonious physical interpretation of these measures is that they aim at representing relative molar concentrations of the RNA in the pool of RNAs analyzed. For instance, the measure of “transcripts per million transcripts” (TPM) is designed to be proportional to the ratio of the molar concentration of the RNA over the sum of the concentrations of all RNA species detected (Wagner et al., 2012):

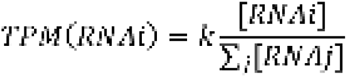

where *k* is a scaling factor and *i* and *j* are indices for genes, and j is running over all genes mapped in that study. Initially, transcript abundance was quantified as RPKM (reads per kilobase per million reads; Mortazavi et al., 2008). RPKM is problematic, however, because it is only consistent for samples with the same distribution of feature lengths (Wagner et al., 2012). Since this cannot be guaranteed when we compare samples from different cell types, organs and species we prefer TPM, which is consistent among samples and has a clear physical interpretation as proportional to the relative molar RNA concentration.

What is lost or not represented in any relative measure is the actual abundance of a RNA species in the cell. Obtaining information about that requires careful analysis of cell size and mRNA concentrations obtained from a given number of cells. Conventional RNA abundance measures, including TPM, RPKM, and FPKM, at best only give information about the relative abundance among the RNA species, rather than their real importance in a particular cell. This needs to be taken into account when we interpret the data from RNAseq studies. Furthermore, transcriptome samples from heterogenous tissues not only reflect transcriptional regulation, but also the relative abundance of cell types in the sample. Thus, heterogeneity of cells within a sample will impact estimates of relative expression, and this must be carefully considered when using transcriptomes to study complex morphological characters.

A consequence of the relative nature of the RNA abundance measures is that their estimates are vulnerable to differential degradation of different RNA species in replicate samples. In Figure 3A we show an example of this problem. We plotted the sqrt(TPM) values from replicate hindlimb samples from H&H stage 29 (Hamburger and Hamilton, 1951) embryonic chickens. When relative expression in biological replicates is compared across all genes one expects a regression line with slope of = 1, and some variation around that line due to biological and experimental variation. In fact, there are subsets of genes that strongly deviate from this expectation in comparisons of some replicates (Fig. 4A), but not between other replicates (Figure 4B). This is caused by differences in RNA quality. If RNA degradation affects all RNA species proportionally, the relative abundances should remain the same and the plot of TPM values should still show the 1:1 relationship expected for replicates. What we see here, however, is a deviation from the 1:1 relationship with some transcripts substantially above the 1:1 diagonal line and some below (Fig. 4A). A likely interpretation is that RNA degradation affects certain RNAs differentially.

**Fig. 4:**
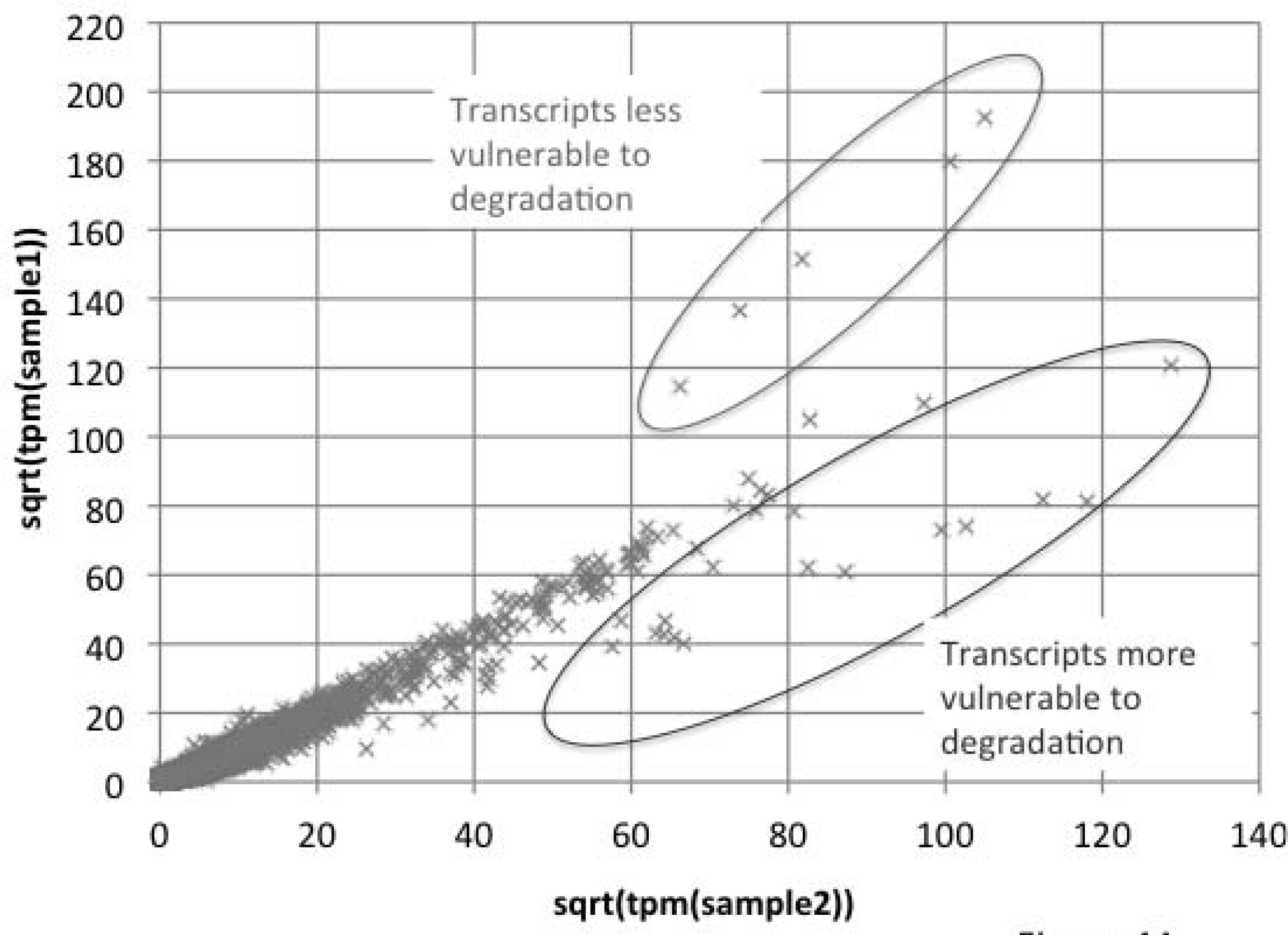
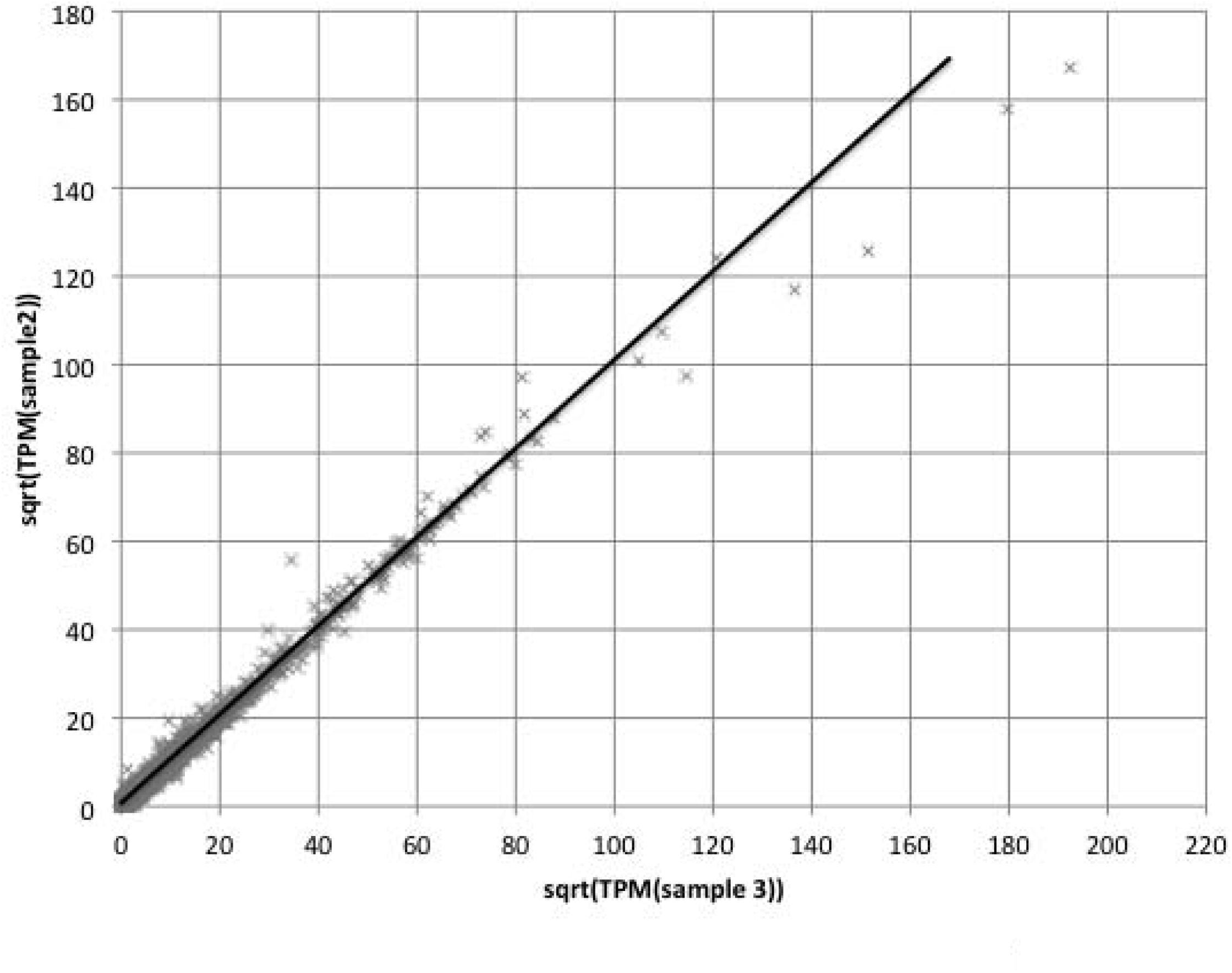
The effect of differential RNA degradation on transcriptome data. A) Comparison of two replicate samples of different quality, where one sample has differentially degraded RNA. B) Comparison of replicate samples with high RNA quality. The relative abundance of corresponding RNA species cluster around the 1:1 regression line with the exception of some highly expressed transcripts. Data from chicken hindlimb buds from Hamburger and Hamilton (H&H; Hamburger and Hamilton, ’51) stage 29.

After obtaining relative RNA expression profiles from our samples, the choice is whether to proceed with the raw relative expression values or to perform a transformation before calculating measures of similarity, such as a correlation matrix, or constructing a tree. We will discuss two directions one may want to go, one where a non-linear transformation is performed, like the log-transformation, prior to analysis, and the other that leads to a discretization of the data into expressed (=1) and non-expressed genes (=0).

The motivation to perform the a non-linear transformation on the TPM or RPKM values comes from the fact that these values can vary over several orders of magnitude, and that similarity measures, like correlations, are vulnerable to the disproportional influence of extreme values in the distribution. The other motivation, in particular when it comes to performing statistical tests to detect differences in expression level, is to eliminate the systematic effect of the mean value on error variance, also called variance stabilization.

The most popular non-linear transformation is the log transformation. In mathematical statistics, this transformation is used when the variance of a random variable is proportional to the square of the mean value of a sample. This is often the case for morphometric variables, i.e. variables that are realized through a multiplicative process, like cell proliferation. It can be shown mathematically that the mean dependency of the variance can be approximately eliminated by a log transformation (Supplemental Material). In the case of transcriptomic data, the problem is that the log transformation, instead of being variance stabilizing, in fact inflates variance for genes with small abundance values. That can be seen in Figure 5A. Another problematic consequence of the log transformation is that the fact that in RNA data abundances of =0 are not uncommon. Of course Log(0)=-infinity, which is not a number one wants to have in one’s data set. Authors often compensate for this by adding an arbitrary small number to TPM or RPKM=0 measures, which in itself shows that the log-transformation is highly problematic, since it forces the investigator to arbitrarily alter the data.

**Fig. 5:**
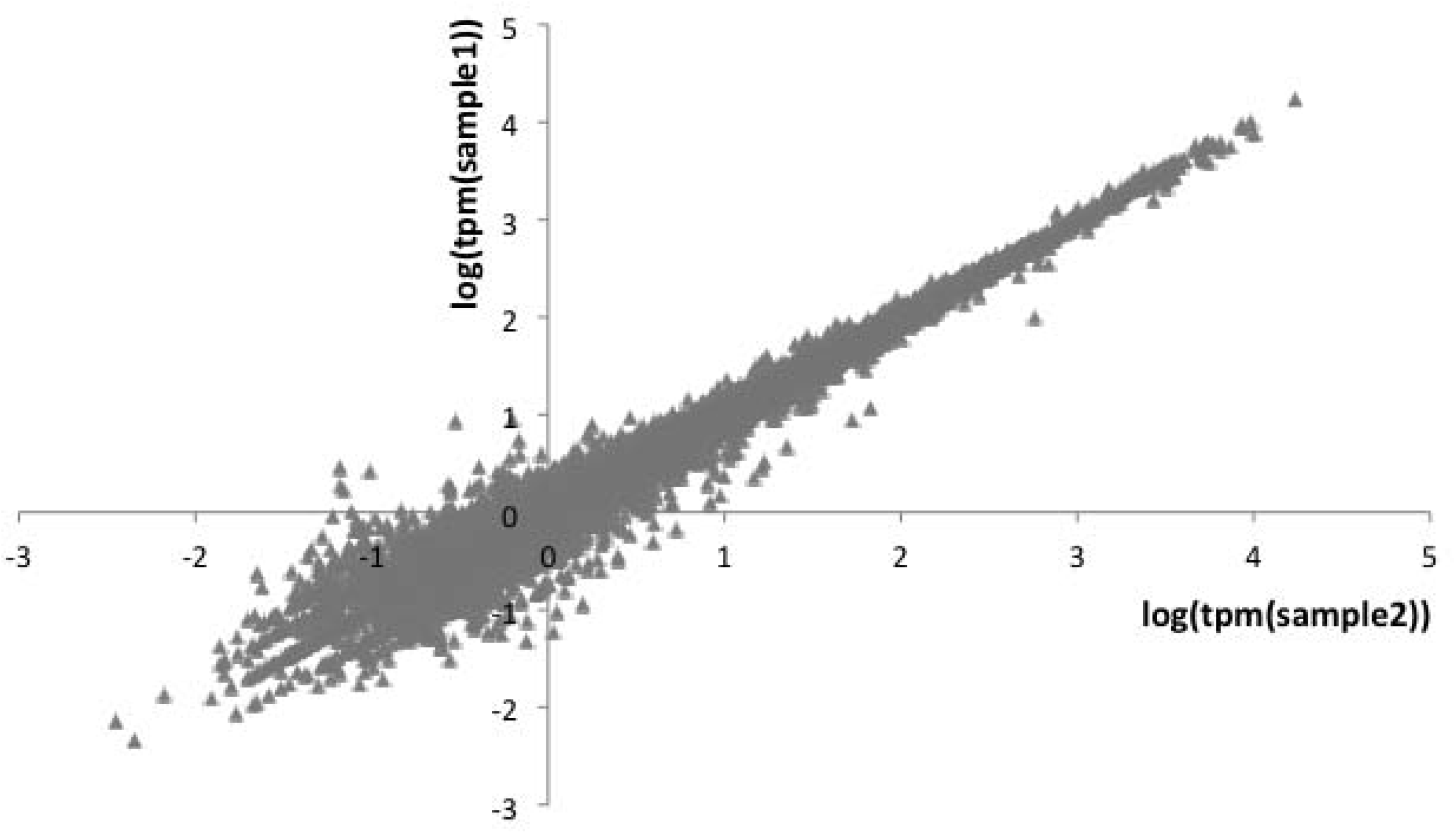
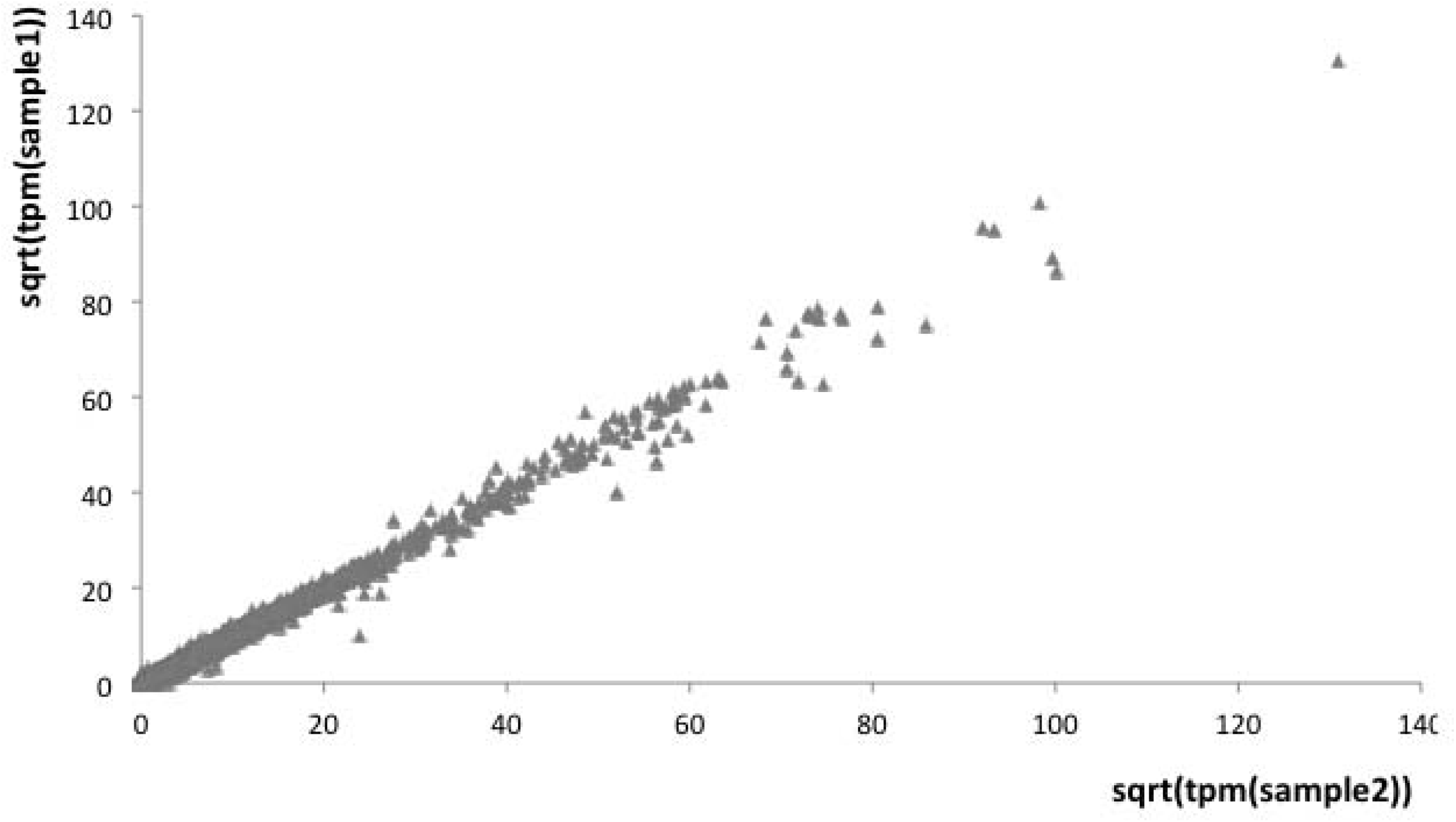
The effect of non-linear transformations on TPM RNA abundance measure. A) Two replicate samples from chick H&H stage 30 forelimb buds after log transformation. Transcripts with zero TPM in one of the samples have been eliminated to avoid the need to plot minus infinity values. Note that the log transformation inflates the variation at low expression levels (tear-drop shape). B) The same two replicate samples as in (A) after square root transformation. Note the complete absence of the tear-drop effect seen after the log transformation. For that reason we recommend square root transformation to be used on transcript abundance data instead of log transformation.

There is, however, an alternative to log-transformation that takes care of these pathologies. To our knowledge this was first pointed out by a blogger who goes by the name *BAMseek* on a blog called *Bridgecrest Bioinformatics* in 2011, <u>http://bridgecrest.blogspot.com/</u>, and advertised on *SEQanswers*. The simplest of these transformations is the square-root transformation. For one, 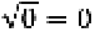 and thus even zero RNA abundances remain in the realm of finite numbers. Even more importantly square-root transformations are actually variance stabilizing for RNA abundance data, and avoid the inflation of variance at low levels (Fig. 5B). From a theoretical point of view square-root transformations are justified when the variance is proportional to the mean value of a sample (Supplemental Material). However, most RNA abundance measures are in fact ratios (see above) and for ratios the appropriate transformation is the arcsine-square-root transformation (see Supplemental Material for a mathematical proof). Since the number of transcripts quantified is usually large, in the tens of thousands, the individual abundances are small compared to the sum of all abundances. For that reason the arcine-square-root transformation can be approximately replaced by the square-root transformation, because the asymptotic slope of the arc-sine function =1 at x=0. *For that reason it is imperative to replace the popular log-transformation by the better square-root transformation* in all applications where transformations are desirable, like tests for differential expression or sample scatter plots and similarity measures among samples (correlations). These correlations can be used for hierarchical clustering and thus is one avenue to obtain classifications that could reflect phylogenetic relationships.

### Expressed/ non-expressed scoring versus fold change comparison

The alternative to using the fully quantified RNA abundance measures in comparing transcriptomes is to classify transcripts into two categories, expressed or not expressed (Wagner et al., 2013). We think that this is an important first step in the analysis of gene expression. There are several reasons for investigating a discretized version of gene expression data. We will discuss these reasons first before we explore how this transformation of RNA abundance data into expressed/non-expressed categorical variables (E/NE) can be done practically.

One reason for considering E/NE description of gene expression profiles is that differences in gene expression can be due to two mechanistically distinct processes: modulation and induction/repression. Modulation happens when a gene that is expressed in both samples but has different expression levels. These changes are usually due to modulations of the activity of upstream regulators, such as the activity of signaling cascades. These changes are appropriately described by fold changes.

In contrast, there are cases where a gene is expressed in one sample but suppressed (“turned off”) in another sample. Expressed and non-expressed genes in eukaryotes often correspond to different states of the chromatin at the locus in the cell. Non-expressed genes are often covered with condensed chromatin, characterized by for instance by H3K27me, i.e. the mono-methylation of the Lysine 27 of histone 3 (Hebenstreit et al., 2011), and the absence of RNA-polymerase II at the promoter (Struhl, 2007). An expressed gene is usually characterized by the presence of Pol-II and H3K4me and often also H3K27ac. Hence, expressed and non-expressed genes are two categorically different states of the gene and differ from cases of gene expression modulation. In many cases in our research we are interested in alternative cell or character identities that are subscribed by qualitative differences in the set of genes expressed, and thus we think that comparing discretized gene expression profiles is biologically more meaningful.

A second reason to compare cell types through their E/NE gene expression profiles is that exact gene expression levels are highly variable and subject to subtle environmental and experimental conditions. In other words, gene expression levels are less likely to be characteristic of cell type identity. We think that as a first step, comparing E/NE gene expression profiles is more reliable rather than taking the full complexities of gene expression modulation on board right away.

Finally, there is a technical reason to prefer E/NE data at this point in time. Ultimately we want the analysis and interpretation of transcriptome data to be based on, and guided by, statistical models of gene expression evolution. It is reasonably straightforward to derive and analyze models with 0/1 binary random variables representing E/NE gene expression data (e.g. Liang et al., 2015). On the other hand, there is no general statistical model that could capture the complex gene expression profiles that covers all differences in gene expression level except generic models for quantitative traits, like the Brownian motion model and models based on the Ornstein-Uhlenbeck process (Felsenstein, ‘88; Hansen and Martins, ‘96). Only a complete gene regulatory network model would be able to do that. To attempt to do that on a genome wide level and across various cell types and species is not realistic at this point.

To be biologically meaningful, however, the transformation of gene expression data into discretized E/NE data the categorization of gene expression states has to be mechanistically justified. It is clear that transcripts with zero reads are likely not expressed. But the problem is that the set of transcripts with zero reads depends not only on the expression level of the gene, but also on the sequencing depth. The more reads that have been reported, the larger the chance that a very low expressed gene is represented with at least one read. Hence, the deeper the sequencing the smaller the number of genes with zero reads mapped to them. Similarly, very small read numbers could be due to leaky transcription rather than indicating an actively transcribed gene.

To obtain a biologically meaningful cutoff value for the classification of genes in expressed and non-expressed there are two approaches, which lead to similar results. One is to use a statistical model (Wagner et al., 2013). The model represents the frequency distribution of genes with different levels of expression, expressed in TPM, for instance. The model consists of the superposition of two statistical distributions, one representing the TPM distribution of genes that are not expressed and another that represents genes that are actively transcribed. The mathematical model is parameterized by regressing the observed frequency distribution of genes in a certain bin of TPM values with the model prediction. The threshold value for the E/NE distinction is then taken as that expression level at which the probability of a gene with a certain expression level or higher to be from the non-expressed category is essentially zero. A comparison of a variety of cell types, suggests that TPM values larger than 2 or 3 are likely from actively transcribed genes (Wagner et al., 2013). This threshold corresponds to roughly 1 RPKM. The second way to categorize genes as expressed or non-expressed is to assay promoter chromatin marks via chip-seq. The first paper attempting this discrimination is by Hebenstreit et al. (2011) and found that the threshold that discriminates between expressed and non-expressed values is about 1 RPKM, which corresponds to the value obtained from the statistical model. Data from our lab on endometrial stromal cells confirms that conclusion (Kin et al., 2015 and Figure 6).

**Fig. 6:**
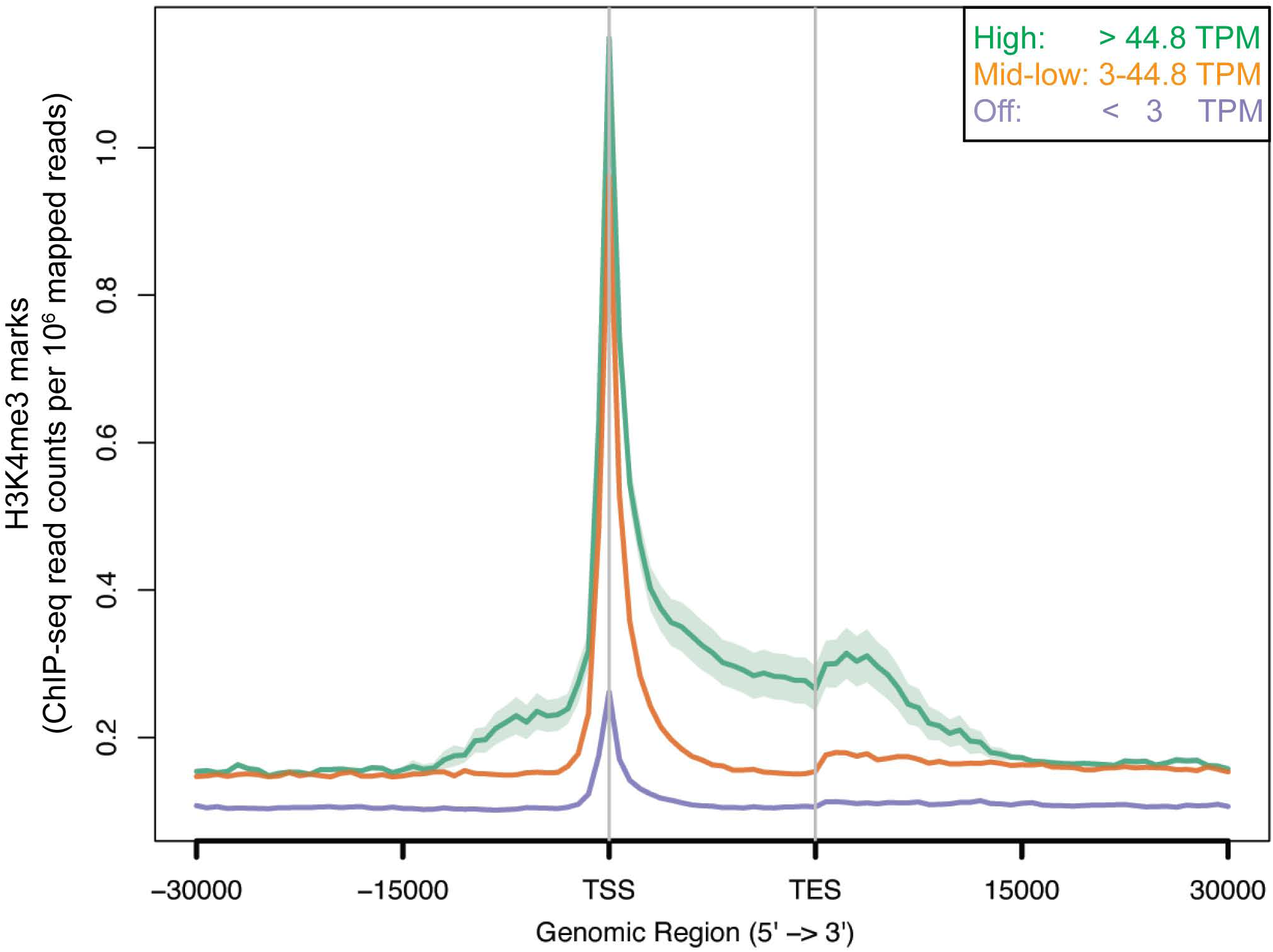
Relationship between average TPM expression levels and amplitude of active promoter mark, H3K4me3. Note that the operationally defined threshold to distinguish between not expressed and expressed genes (3tpm) corresponds in a large difference in the level of H3K4me3 active prompter mark in a ChIPseq dataset, whereas the difference between moderately expressed genes (3 to 45 tpm) and highly expressed genes is comparatively small (Figure 2 from Kin et al, 2015).

When one applies an operational threshold of say ≥2tpm one will have a large number of genes that are in the gray-zone around the threshold. To deal with that one could apply one sample t-tests to assess whether a gene expression level in a sample is significantly larger than the threshold value, although this may be useful only when 3 or more replicates are available. In many applications one can apply some operational “no-man’s land” between expression levels that are classified as expressed and non-expressed. For instance on may say a gene is expressed if it has a tpm>2x the expression threshold of 2tpm (i.e. ≥4tpm) and not expessed if tpm is < 1, and genes that fall inbetween these two thresholds are considered as ambiguous. According to our experience, the exact threshold levels used have little effect on the results,.

### Challenges of inter-species comparisons

There are two problems when comparing transcriptomes from different species. The one is that TPM values depend on the set of genes mapped in each species. This is obvious given that TPM of a particular gene is the number of transcript sequences in a sample divided by the number of sequenced and mapped transcripts in that sample (see above; Wagner et al., 2012). If we have data from two species, A and B, and the number of annotated genes in A and B are different, then the TPM values for genes mapped in both species will be systematically different, even when the expression levels are exactly the same, because the denominator of the TPM calculation, the total number of transcripts sequenced and mapped, is different. We will address this problem below. The other difficulty is the fact that the quality of gene annotation can be different between the genomes of the species compared. Mapping to different gene models can affect the estimated gene expression level if the profile of mapping to parts of the gene model is non-uniform. A possible solution of this problem is to construct a minimal gene model consisting of orthologous sequences that are shared among all species compared (Pankey et al., 2014).

Above we noted that TPM estimates have to differ in samples from two species if the number of annotated genes is different. There is, however, a simple approximate method to rescale these values to make TPM values comparable among species. Let us consider two species A and B. One of the two species, say A, is a model species with a more completely annotated genome than B. Let us further assume that the genes annotated in B are a proper subset of the genes annotated in A, *G*_*B*_ ⊂ *G*_*A*_, and ‖*G*_*B*_‖ = *N*_1_ as well as ‖*G*_*A*_ ‖ = *N*_2_, with N_1_< N_2_. The number of transcripts sequenced of gene *i* is

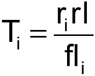

with *r*_*i*_ the number of reads mapped, *rl* the length of the reads and *fl*_*i*_ the feature length of gene *i*. Since the read length is the same across all genes within a sample it cancels from the calculation of any ratio and we can simply proceed with a simpler measure of transcript abundance in a sample: 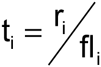. Note that *t*_*i*_ still depends on the sequencing depth, i.e. the number of reads, and thus is a sample specific value.

In order to rescale the TPM values from two species with different numbers of annotated genes we have to make a crucial assumption:

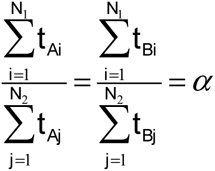

This assumption means that the relative transcript abundances in the two gene sets *G*_*A*_ ∩ *G*_*B*_ (the overlapping set) and *G*_*A*_¬*G*_*B*_ (the set not mapped in B but assumed to be there) are approximately the same in the two species and samples. Now let us compare the TPM values in the original quantification of samples from species B, *tpm*_*Bi*_, and the TPM rescaled to the same larger gene set *G*_*A*_: 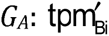.

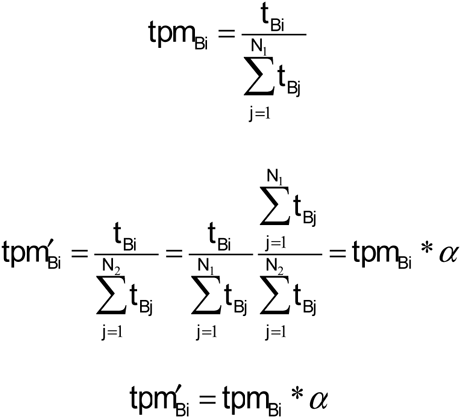

In words: to rescale the TPM values in species B we only need to calculate the α ratio from the samples from the reference species A and multiply the TPM values from species B with that α-ratio. From a practical point of view the α-ratio is easily calculated by

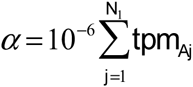

since per definition 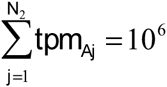 and the denominator of the TPM cancels when calculating the ratio.

## Conclusions

The comparative analysis of transcriptome data from different morphological characters provides insights into various aspects of character evolution. The factors influencing the structure of morphological character trees includes the evolutionary relationships among the characters or cell types compared, the phylogenetic relationships among the species from which the characters were sampled, and the degree of variational individuation of the characters. In some cases, particularly among closely related characters, developmental origin may also shape the character tree. For that reason, the tree structure inferred from transcriptome data has to be carefully interpreted. Any interpretation should take into account information from independent sources, including the species phylogeny and the likely phylogenetic age of the characters. Ideally, transcriptomes from different morphological characters can be sampled from multiple species with well-established relationships. For investigation of novel characters, potential serial homologues should be sampled from lineages that bracket the origination of the character under investigation. However, if carefully evaluated, the comparative analysis of transcriptome data can be a powerful tool for investigating the evolution of morphological characters and their underlying gene regulatory networks. Further, this approach offers a new opportunity to test hypotheses regarding the evolution of morphological disparity, first described at the morphological level, but now understandable at the genetic level and in causal detail.

## Acknowledgments

We would like to thank Todd Oakley, Thomas Stewart, and two anonymous reviewers for their thoughtful and helpful reviews of the manuscript. The authors also want to thanks Cong Liang for help in deriving the between sample scaling procedure described here as well as acknowledge collaboration on the topic of the species signal that will be presented in a forthcoming paper with Ms Liang as first author.

Research in the Wagner lab has been supported by a research grant from Foundational Questions in Evolutionary Biology (#R11636) for work on limb transcriptomics, a research grant from the John Templeton Foundation (#54860) to investigate cell type evolution, and by the Yale University science development fund. The authors also acknowledge funding from the Dillon and Mary Ripley Graduate Fellowship Fund at Yale University (JMM) and the National Science Foundation Graduate Research Fellowship under grant No. DGE-1122492 (JMM).

## Supplementary Material

The results presented here are most likely not original, but have not been presented in any of the statistics textbooks the authors have examined. One of us (GPW) has derived these results for his statistics course and found it highly useful to students to understand why certain transformations work under what circumstances.

The results below use the following fact: let z=f(x) and let the curvature of f(x) at the mean of M(x) be smaller than the standard deviation of x then

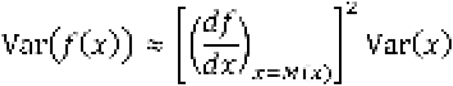

**Result 1:** if the variance of a random variable *x* is proportional to the square of the mean, *Var(x)= k M(x)*^2^, then the variance of the log-transformed variable z=log_a_(x) is approximately invariant with respect to mean of z.

**Proof:**

Since

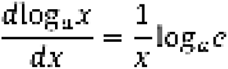

we have

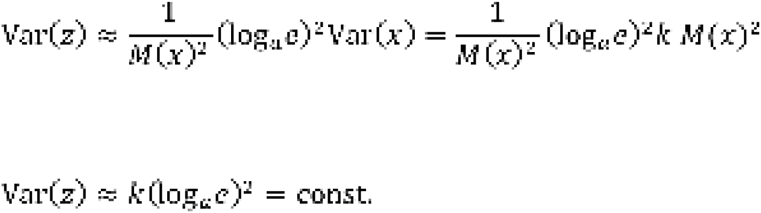

**Result 2**: if the variance of a random variable *x* is proportional to the mean, Var(*x*)=*k* M(*x*)^2^, then the variance of the square-root transformed variable 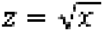 is approximately invariant with respect to mean of *z*.

**Proof:**

Since

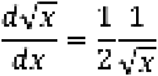

and thus we have

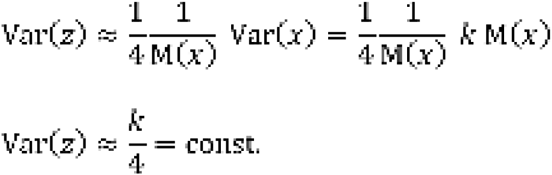

**Result 3:** let us consider a random variable *x* where Var(*x*)=k*M(*x*)[1-M(*x*)], then the variance of 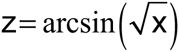 is approximately invariant of M(z).

Note that this situation is for instance the case for ratios of counts, like the relative frequency of a Bernulli variable. For that reason it is a good model for relative RNA abundance measures.

**Proof:**

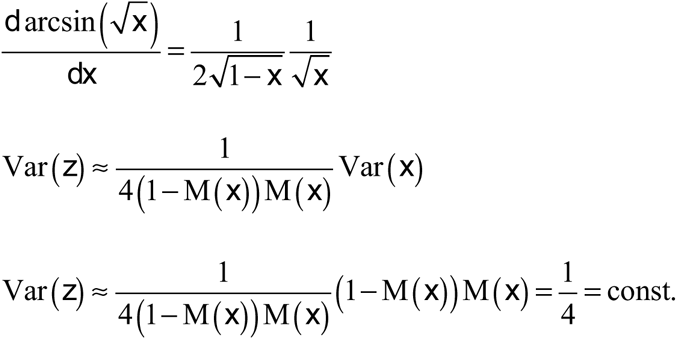

**Result 3**: let us consider a random variable *x* where Var(*x*)=k*M(*x*)[1-M(*x*)], then the variance of 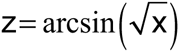 is approximately invariant of M(z).

Note that this situation is for instance the case for ratios of counts, like the relative frequency of a Bernulli variable. For that reason it is a good model for relative RNA abundance measures.

**Proof:**

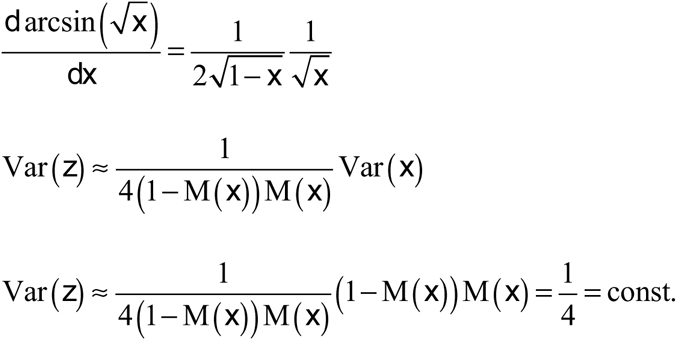

## Footnotes

1 http://www.molecularecologist.com/2015/05/another-lesson-in-genomics-experimental-design-and-avoiding-batch-effects/#more-6265

